# Perception of self- and externally-generated visual stimuli: Evidence from EEG and behaviour

**DOI:** 10.1101/2022.05.30.492531

**Authors:** Edward Ody, Benjamin Straube, Yifei He, Tilo Kircher

## Abstract

Efference copy-based forward model mechanisms may help us to distinguish between self- and externally-generated sensory consequences. Previous studies have shown that neural activity and perception associated with self-initiated sensory consequences are reduced (sensory suppression). For example, event-related potentials (ERPs) elicited by tones that follow a button press are reduced in amplitude relative to ERPs elicited by passively attended tones. However, previous EEG studies investigating visual stimuli in this context are rare, provide inconclusive results, and lack adequate control conditions with passive movements. Furthermore, although self-initiation is known to modulate behavioural responses, it is not known whether differences in the amplitude of ERPs also reflect differences in perception of sensory outcomes. In this study, we presented to participants visual stimuli consisting of grey discs following either active button presses, or passive button presses, in which an electromagnet moved the participant’s finger. Two discs presented visually 500-1250 ms apart followed each button press, and participants judged which of the two was more intense. Early components of the primary visual response (N1 and P2) over the occipital electrodes were suppressed in the active condition. Interestingly, suppression in the intensity judgement task was only correlated with suppression of the visual P2 component. These data support the notion of efference copy-based forward model predictions in the visual sensory modality, but especially later processes (P2) seem to be perceptually relevant. Taken together, the results challenge the assumption that N1 differences reflect perceptual suppression and emphasise the relevance of the P2 ERP component.

## 1. Introduction

Neural and behavioural responses elicited by self-generated sensations are attenuated compared to those elicited by externally-generated sensations (for example, Aliu et al., 2009; Blakemore et al., 1998; Uhlmann et al., 2020). According to forward model theories of sensorimotor control (Wolpert, 1997; Wolpert & Ghahramani, 2000), when an action is performed, a copy of the motor command for that action is created (known as the *efference copy*; von Holst & Mittelstaedt, 1950), which represents a prediction for the outcome of that action. The efference copy is then compared with the actual sensory feedback. If the two match, the perception of that feedback, along with cortical activity associated with it (e.g. Blakemore et al., 1998; Shergill et al. 2013) is attenuated or cancelled out, resulting in the perception of those stimuli as less intense. In addition to helping us distinguish between self- and externally-generated sensory action consequences, this mechanism may function to increase efficiency by preventing the allocation of attention to irrelevant self-produced sensations (Pynn & DeSouza, 2013).

The neural underpinnings of this phenomenon have been extensively investigated using event-related potentials (ERPs). In the auditory sensory modality, N1 and P2 ERPs elicited by tones following a button press are reduced in amplitude relative to identical tones which are passively attended (e.g. Bäß et al., 2008; Martikainen et al., 2005; Mifsud et al. 2016; SanMiguel et al., 2013). This effect is sometimes called sensory suppression and has been reported in numerous auditory domain studies. In the visual domain, however, relatively few studies have investigated sensory suppression of self-generated action consequences and the results of these have been less consistent. Two previous studies (Mifsud et al., 2018; Schafer & Marcus, 1973) reported N1 suppression for self-generated flashes over the vertex electrodes (though Schafer & Marcus reported no significant differences over occipital electrodes). Similarly, Gentsch & Schütz-Bosbach (2011) reported N1 suppression of self-generated arrow shape stimuli in fronto-central, central, and central-parietal electrodes. Csifcsák et al. (2019) found suppression of occipital C1 for both self-generated chequerboard and hand stimuli. However, the amplitude of the later P1 component was larger for self-than externally-generated stimuli. Similarly, Hughes and Waszak (2011) found a significantly increased P1 amplitude for self-generated chequerboard stimuli. However, the later period, between 150 and 350ms after stimulus onset was reduced for self-generated stimuli. Finally, Mifsud et al. (2016) observed an enhanced N145 response over occipital electrodes for self-generated pattern reversals. These results are therefore inconsistent in providing evidence for an efference copy-based forward model mechanism in the visual domain.

The majority of previous sensory suppression EEG experiments have shared a similar design, called the contingent paradigm (Horváth, 2015). In the active (self-generated) condition, participants press a button which triggers a stimulus. In the passive (externally-generated) condition, the stimuli are most commonly presented with the same temporal sequence and participants are asked to simply attend to them. Typically, a motor control condition is also included. In this condition, participants press the button but no stimulus follows. The ERP waveform from this condition is subtracted from that of the active condition. It is assumed that this will remove any button press-related activity from the active condition and that the active and passive conditions will then differ only in the presence of the efference copy.

This assumption has been questioned on the basis that there could be confounding differences between the active and passive conditions (Horváth, 2015). In the active condition, participants are engaged in an additional task (pressing the button) and obtain additional sensory information related to the button press compared to the passive condition, where they are asked to passively attend to the stimuli. Attention is known to increase the amplitude of N1 potentials (e.g. Hillyard et al., 1973; Hillyard, 1981; Nobre, 2001). In the contingent paradigm, it is assumed that participants will focus attention on the stimuli in the passive condition, either spontaneously to retain focus or simply because this is what they are asked to do, leading to an increase in N1 amplitude. However, in the active condition, attention is instead focussed on the action at the time it is performed. This could lead to an apparent attenuation effect in the active condition. Tones randomly coinciding with a button press have been shown to elicit attenuated N1 and P2 responses (Horváth et al., 2012), suggesting that the mere close temporal proximity between button press and tone is sufficient to attenuate sensory processing.

Secondly, it has been argued that there are inherent differences in temporal prediction between the active and passive conditions in the contingent paradigm that could affect the results (Horváth, 2015). Stimulus timing may be more predictable in the active condition than the passive condition. This is because participants are able to prepare for the stimulus based on the tactile and proprioceptive information of the initiating button press. In the passive condition, no such warning of the stimulus occurs. Previous studies have shown mixed evidence to support this hypothesis. Sowman et al. (2012) showed that N1 for self-initiated and externally cued sounds did not significantly differ in amplitude. Bäß et al. (2008) compared self-initiated sounds with temporally predictable and unpredictable stimulus delays. They found that the self-initiated sounds with predictable delays led to greater sensory suppression. These studies suggest that temporal predictability plays a role in the processing of self-initiated sounds. However, Lange (2011) has shown that N1 elicited by sounds generated by a keypress have a lower amplitude compared to sounds following a visual cue consisting of a grey square. Klaffehn et al. (2019) similarly showed that when sounds were made predictable using a continuously filling up bar, N1 was still smaller following self-compared to externally-generated sounds. These results conversely suggest that movements do contain a unique predictive component that is separate from the inherent temporal prediction that they allow for. Nevertheless, temporal prediction is a factor that should be controlled in an optimum design.

As previously mentioned, according to forward model theories of sensorimotor control, not only neural activity related to self-generated sensory action consequences is reduced but also perception of those stimuli on the behavioural level. For example, tickling sensations are reduced when one tries to tickle oneself, relative to when the tickling is done by another person or a machine (Blakemore et al., 1998, 2000). In auditory (Desantis et al, 2012; Sato 2009; Weiss et al., 2011a, 2011b) and visual (Lubinus et al., 2021) intensity judgement tasks, self-generated stimuli are perceived as less intense than externally-generated stimuli. It is known that the amplitude of the auditory N1 component is related to stimulus intensity, with louder sounds producing larger N1 amplitudes (Kaskey et al., 1980). Therefore, for example, as reduced N1 amplitude elicited by self-generated tones have often been interpreted as reflecting a reduced intensity perception of those tones, no previous studies, to our knowledge, have directly tested this correlation between neural and behavioural data.

The present study had three aims. Firstly, we wanted to investigate sensory suppression of visual action consequences with optimal passive control conditions. To allow us to compare results in our novel design with previous experiments, we also included a comparable auditory condition. To our knowledge, only one prior study (Mifsud et al., 2016) has investigated both modalities using a within-subject design. Secondly, we aimed to test the link between EEG-measured neural activity and behaviour by including an intensity judgement task. Lastly, but most importantly, we wanted to create an experimental design in which we could optimally control attention, temporal predictability and tactile/proprioceptive feedback.

To this end, we employed a custom-built button box (*passive movement device*; PMD). The device can be operated like an ordinary button or produce an involuntary movement, pulling the finger down by means of an electromagnet (for details see section 1.2). We have investigated action-related predictive processes using this and other similar PMDs in our previous behavioural (Arikan et al., 2017; van Kemenade et al., 2016), fMRI (Arikan et al., 2019; Kavroulakis, et al., 2021; Pazen et al., 2020; Schmitter et al, 2021; Straube et al., 2017; Uhlmann et al., 2020; Uhlmann et al., 2021; van Kemenade et al., 2019) and tDCS (Straube et al., 2020) studies. However, to the authors’ knowledge, no previous EEG sensory suppression studies have implemented a PMD. Using this device allowed for more closely matched temporal prediction of the stimuli between the active and passive conditions because the button press acted as a cue in both. Furthermore, tactile feedback from the button was present in both conditions. Finally, task differences between the two conditions were reduced, as a button press was present in both.

We expected that the typical results of auditory sensory suppression (reduced N1 and P2 for self-generated tones) would be present for the visual modality, too. We hypothesised that the amplitude of early components of the primary visual response would also be reduced for self-generated visual stimuli. Regarding behaviour, we hypothesised that active stimuli would have lower points of subjective equality than passive stimuli (indicating that they were perceived as less intense).Finally, we hypothesised that suppression in the intensity judgement task and suppression of the elicited neural activity would be correlated.

## 2. Method

### 2.1 Participants

29 participants took part in the study, of which 19 were female. Ages ranged between 19 and 31 (M = 23.7, SD = 3.35). Participants were recruited through a university mailing list and were therefore mostly students or staff at the University of Marburg. Participants received a €30 inconvenience allowance for taking part. The study protocol was approved by the local ethics committee in accordance with the Declaration of Helsinki (except for pre-registration; World Medical Association, 2013), and all participants provided written informed consent. By self report, all participants were right-handed, had normal or corrected-to-normal hearing and vision, no history of mental illness, no history of drug or alcohol abuse, no history of serious brain injury and no first-degree relatives with Schizophrenia. All participants were naïve to the purpose of the experiment. Due to a technical error with the presentation software, one participant’s behavioural data were missing. This participant was included in EEG-only analyses but excluded from any analyses including behavioural data. A second participant was missing both data types and was therefore removed from both analyses. In total, 27 participant’s data were included in behavioural analyses and 28 in EEG analyses. To verify the results, we also completed the same EEG analyses with only the 27 participants that were included in the behavioural analyses. This has led to largely comparable EEG results (see Appendix).

### 2.2 Task and Procedure

Figure 1 illustrates the task procedure and PMD. The experiment took place in a semi-darkened room. Participants sat at a comfortable distance behind a 19” 60Hz computer monitor. They placed their right hand on a button pad with the right index finger resting on the button and attached securely to it with a piece of elastic fabric. The participant’s left hand was placed on the computer keyboard, on which the response keys (‘v’ and ‘n’) were marked. Pink noise was played through headphones throughout the experiment to mask the sound of the button press. Participants first completed a training session to familiarise them with the task. This consisted of 5 blocks of 5 trials each. The EEG cap was then fitted prior to starting the main experimental and control blocks.

**Figure 1.**
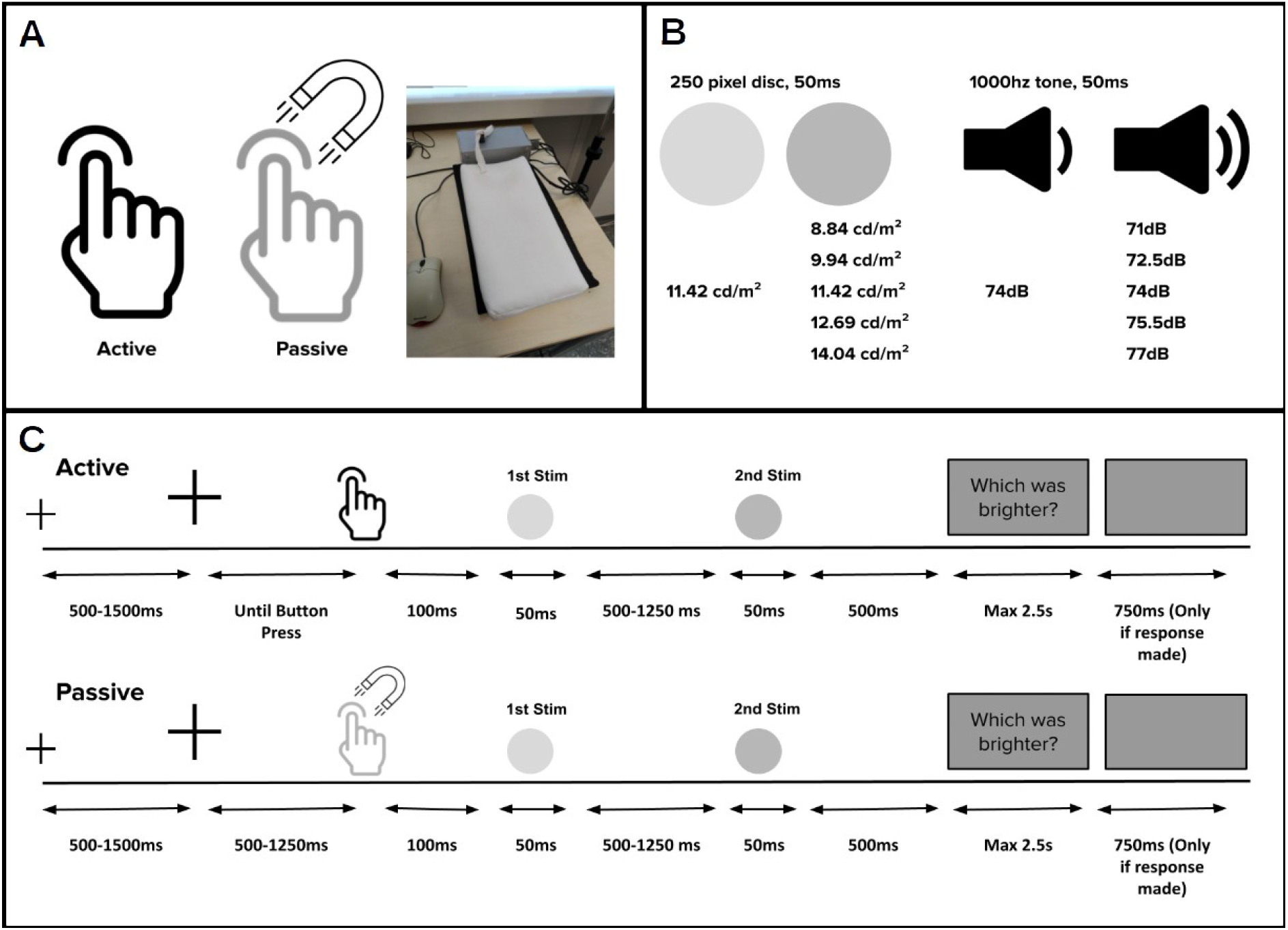
Schematic illustration of the task. A) The PMD. Active button presses were made by the participant. Passive button presses moved the finger down using an electromagnet. B) Stimulus details. Participants judged which of the two stimuli presented in each trial was brighter/louder. C) Trial structure (with visual stimuli are shown for illustration purposes). Control conditions were identical, except that the delay between button press and first stimulus was extended to 1000ms.

Participants completed visual and auditory intensity judgement tasks in which they had to decide which of two successive stimuli was brighter or louder. The first stimulus was triggered by the PMD, operated with the participant’s right index finger. The button was either pressed actively by the participant (active condition) or would move the participant’s finger by means of an electromagnet (passive condition). The task consisted of judging which of two grey discs was brighter (visual condition), or which of two tones was louder (auditory condition). The first stimulus (stimulus of interest) was always displayed at a fixed intensity, while the second (comparison stimulus) varied in intensity. Participants indicated which stimulus had the higher intensity by pressing ‘v’ (first) or ‘n’ (second) on the keyboard.

The experiment was divided into 2 experimental blocks and 2 control blocks. Each main block contained 200 trials which were further divided into active and passive mini blocks containing 25 trials per mini block (making a total of 4 active and 4 passive mini blocks per main block). Participants always completed both of the auditory blocks together and both of the visual blocks together. Counterbalancing was achieved by varying the order of presentation (visual or auditory first) and the starting action in each block (active or passive movement). The data were collected during a session which also included multimodal blocks which were not the focus of the present study. These blocks were interspersed with the described blocks. All visual blocks and all auditory blocks were always presented together. However, the order of unimodal and multimodal blocks, the order of auditory and visual blocks and the order of active and passive trials within the blocks was counterbalanced. 50% of participants started with visual and 50% started with auditory. 50% started with unimodal while 50% started with multimodal. 15 participants started with active trials while 13 started with passive.

At the start of each trial, a black fixation cross was presented. After a randomly selected duration between 500 and 1500ms (in 250ms steps), the cross enlarged. In active trials, this cue indicated that participants could press the button at their own pace. In passive trials, it indicated that the passive button would shortly pull the finger down. In these trials, the button was activated after a randomly selected interval between 500 and 1250ms (in 83ms intervals). Following each button press, regardless of active or passive, there was a 100ms delay, after which the first stimulus (stimulus of interest) was presented. This delay was included to match temporal prediction between active and passive trials, so that even though participants could not predict when their finger would move in the passive condition, they would be able to predict when the stimulus would appear. It could be potentially argued that introducing a delay between the button press and stimulus could reduce sensory attenuation. However, Aliu et al. (2009) have shown that attenuation of auditory action consequences does occur at delays of 100ms. A second stimulus (comparison stimulus) was presented after a randomly selected interval with variable duration between 500 and 1250ms (in 250ms intervals). After a 500ms pause, a question was shown. ‘Welcher war heller?’ (‘which was brighter?’, in German) was shown on visual trials and ‘Welcher war lauter?’ (‘which was louder?’) on auditory trials. Participants pressed ‘v’ on the keyboard to indicate ‘first stimulus’ and ‘n’ to indicate ‘second stimulus’. Making a response triggered a 750ms inter-trial interval. If no response was made after 2500ms, the next trial started automatically. In auditory trials, the fixation cross remained on screen during stimulus presentation and disappeared only when the question was presented. In visual trials, the fixation cross disappeared while the stimuli were displayed but remained on screen during the inter-stimulus interval and between the stimulus and question. Comparison stimuli were presented in a pseudorandom order where the same stimulus level was not shown consecutively more than twice. A diagram of the trial structure is shown in Figure 1C.

The control conditions consisted of 60 active and 60 passive trials. Here, the fixation cross, button press, visual and auditory stimuli, and inter-trial interval were identical to the experimental conditions. However, after the button press there was a 1000ms delay before the stimuli and question were presented. In the typical paradigm, participants simply press the button, with no stimulus presented (Horváth, 2015). However, we felt that this method would allow us to capture the button press activity while keeping participants engaged in the same way as in the experimental conditions. In the control condition, active and passive trials were presented in mini blocks of 15 trials each. The first 60 trials had the visual task while the second 60 had the auditory task.

### 2.3 Stimuli

Stimuli were presented with Psychtoolbox (V 3.0.12) running on Octave (V 4.0.0) in Linux. Auditory stimuli consisted of a 1000hz tone. The first tone was always presented at 74dB, whereas the second had a loudness of 71, 72.5, 74, 75.5 or 77dB. Visual stimuli consisted of a solid 250-pixel disc. The first disc was always presented at a luminance of 11.42 cd/m², whereas the second had a luminance of 8.84, 9.94, 11.42, 12.69 or 14.04 cd/m². Stimuli were presented for 50ms. Luminance measurements were performed using an i1Display Pro photometer (X-Rite Pantone, Grand Rapids, USA). Volume measurements were performed using an RS-95 decibel metre (RS Components Ltd). The stimuli were presented on a fixed grey background with luminance of 3.40cd/m².

### 2.4 EEG Data Acquisition

EEG was continuously recorded at a sampling rate of 500Hz from 32 active Ag/AgCl electrodes (Fp1/2, F7/8, F3/4, Fz, FT9/10, FC5/6, FC1/2, T7/8, C3/4, Cz, TP9/10, CP5/6, CP1/2, P7/8, P3/4, Pz, O1/2 and Oz). The EEG was referenced online to the electrode location Fcz and the ground electrode was placed on the forehead. EEG were filtered online between Impedances were kept at 25 kO or below. The signal was amplified by a BrainVision amplifier and recorded with BrainVision Recorder (Brain Products GmbH, Germany). Electrodes were mounted in an elastic cap (actiCAP, Brain Products GmbH, Germany) according to the international 10–20 system.

### 2.5 EEG Preprocessing

Data preprocessing was implemented firstly with the EEGLAB toolbox (Delorme & Makeig, 2004). Raw continuous EEG were firstly downsampled to 250Hz and then high-pass filtered at 0.5Hz. Line noise was removed using cleanLineNoise from the PREP pipeline (Bigdely-Shamlo et al., 2015). Bad channels were identified using artefact subspace reconstruction (Chang et al., 2018) and interpolated. Resulting EEG were rereferenced to the average of the two mastoid electrodes, T7/8. Then, automatic ICA-based artefact detection and rejection was implemented with AMICA (Palmer et al., 2012). Some active trials had unusually short button press latencies. This could have resulted from the button being partially pressed down at the start of the trial, causing it to register a press and trigger the stimuli immediately. Trials with button press latencies shorter than 100 ms (200 trials, 1.49%) and trials in which no response (92 trials, 0.68%) were removed from all subsequent analyses. 1 trial had both an invalid button press latency and a missed response and was also removed. Data were then low-pass filtered at 20 Hz, and an additional artefact identification and rejection procedure was implemented with the ft_artifact_zvalue (z = 12) function from the Fieldtrip toolbox (Oostenveld et al., 2011). This resulted in removal of a further 12.1% (1611) of the remaining trials. Further analyses were completed with the Fieldtrip toolbox and custom routines in MATLAB (R2020a Mathworks, Sherborn, Massachusetts).

### 2.6 Intensity Judgement Task

Performance in the intensity judgement task was assessed by examining the proportion of trials in which the stimulus of interest (first stimulus) was perceived as darker/quieter. Since the luminance/loudness of the stimulus of interest was held constant throughout the experiment, any change in perception of brightness or loudness would reflect purely perceptual differences. The proportion of trials in which participants answered ‘second stimulus brighter/louder’ were calculated for each participant, condition and stimulus level. Logistic psychometric functions were fitted using Psignifit 4 (Schütt et al., 2015) for MATLAB, implementing a maximum-likelihood estimation, as described in Wichmann and Hill (2001). Thresholds were derived for each participant and condition. The threshold reflects the intensity value at which the participant judged the stimulus of interest and comparison stimulus to be the same intensity in 50% of the trials (point of subjective equality; PSE). When comparing thresholds across conditions, a shift towards lower intensities indicates that the comparison stimulus was judged as brighter or louder more often than the stimulus of interest (for a comparable approach see, Lubinus et al. 2021). We expected perception of self-generated stimuli to be attenuated and therefore our hypothesis was that thresholds would be significantly lower in the active compared to the passive condition. Individual psychometric functions were visually inspected. Participants showing grossly nonconforming functions in either visual or auditory conditions were removed from all analyses (two participants). This resulted in removal of two participants from all subsequent analyses. For statistical analysis, the auditory and visual thresholds were entered separately into 2-tailed paired samples t-tests.

### 2.7 ERP- N1/P2

Data were segmented from 500ms before to 1000ms after the button press. The data were then baseline-corrected to the 200ms interval preceding button press. Next, the mean activity per participant, channel and time point was calculated for each condition. The activity in the active and passive control conditions was then subtracted from the equivalent active/passive auditory and visual conditions. This method has been employed in previous experiments focussing on both auditory (e.g. Bäß, e al, 2008; Whitford et al., 2011) and visual (Csifcsák et al., 2019; Mifsud et al., 2016; Mifsud et al., 2018) ERPs. Finally, data were baseline-corrected to the 100ms interval between the button press and stimulus. For visual conditions, the ERP was averaged across channels O1, O2 and Oz and for auditory conditions across C3, C4 and Cz.

For statistical analysis, peak values were extracted by identifying the minimum (N1) or maximum (P2) value of the ERP across all conditions. A 24ms time window was defined around this centre value. For visual ERPs, this procedure resulted in time windows of 84-108ms for N1 and 132-156ms for P2 and for auditory ERPs 80-104ms for N1 and 144-168ms for P2. The activity in these time windows was averaged together to produce a single value per condition, per participant. Comparable methods have been used in similar previous studies (e.g. Mifsud et al., 2016; Saupe et al., 2013). Active and passive means were compared with a paired samples t-test.

### 2.8 Correlation between ERP and behaviour

To investigate whether perceptual suppression (lower perceived intensity) of self-generated visual and auditory action consequences was related to suppression of the resulting neural activity, we performed correlation analyses. Pearson correlation coefficients were calculated to test the correlation between suppression in the behavioural task and suppression of both N1 and P2, separately for the auditory and visual conditions. Suppression in the behavioural task was calculated by subtracting active thresholds from passive thresholds (passive-active; cf. above). Due to the difference in polarity, N1 suppression was calculated by subtracting the passive condition from the active condition (active-passive) while P2 suppression was calculated by subtracting the active condition from the passive condition (passive-active). In all of these calculations, positive values indicate suppression of self-generated sensory action consequences (lower perceived intensity and lower N1/P2 amplitude in active vs. passive conditions) while negative values indicate enhancement (higher perceived intensity and higher N1/P2 amplitude in active vs. passive conditions). A positive correlation would indicate that participants who showed stronger suppression in the behavioural task, also showed stronger neural suppression. Correlation analyses were secondary to the main comparison of N1 and P2 amplitude between active and passive conditions and were performed only when the expected amplitude differences were observed. Results were correct regarding the number of performed tests (Bonferroni-corrected alpha level of .025).

## 3. Results

### 3.1 Intensity Judgement Task

There was no significant difference in thresholds between the active and passive conditions for the visual condition (*t_24_* = .83, *p* = .41, *d* = .17, CI = [−0.136, 0.319]) or auditory condition (*t_24_* = .92, *p* = .37, *d* = .18, CI = [−0.086, 0.223]).

### 3.2 EEG- N1/P2

For the visual condition (Figure 2), there was a significant (*t_25_* = 4.3, *p* < .001, *d* = .84, CI = [0.749, 2.124]) effect of action type, with more negative N1 peak values for passive (mean = −.99) than active (mean = .44) movements, indicating N1 suppression. For the P2 peak values, the effect of active and passive movement types was also significant (*t_25_*= −4.14, *p* < .001, *d* = −.81, CI = [−2.118, 0.71]) with more positive values for passive (mean = 8.57) than active (mean = 7.16) movements, indicating P2 suppression.

**Figure 2.**
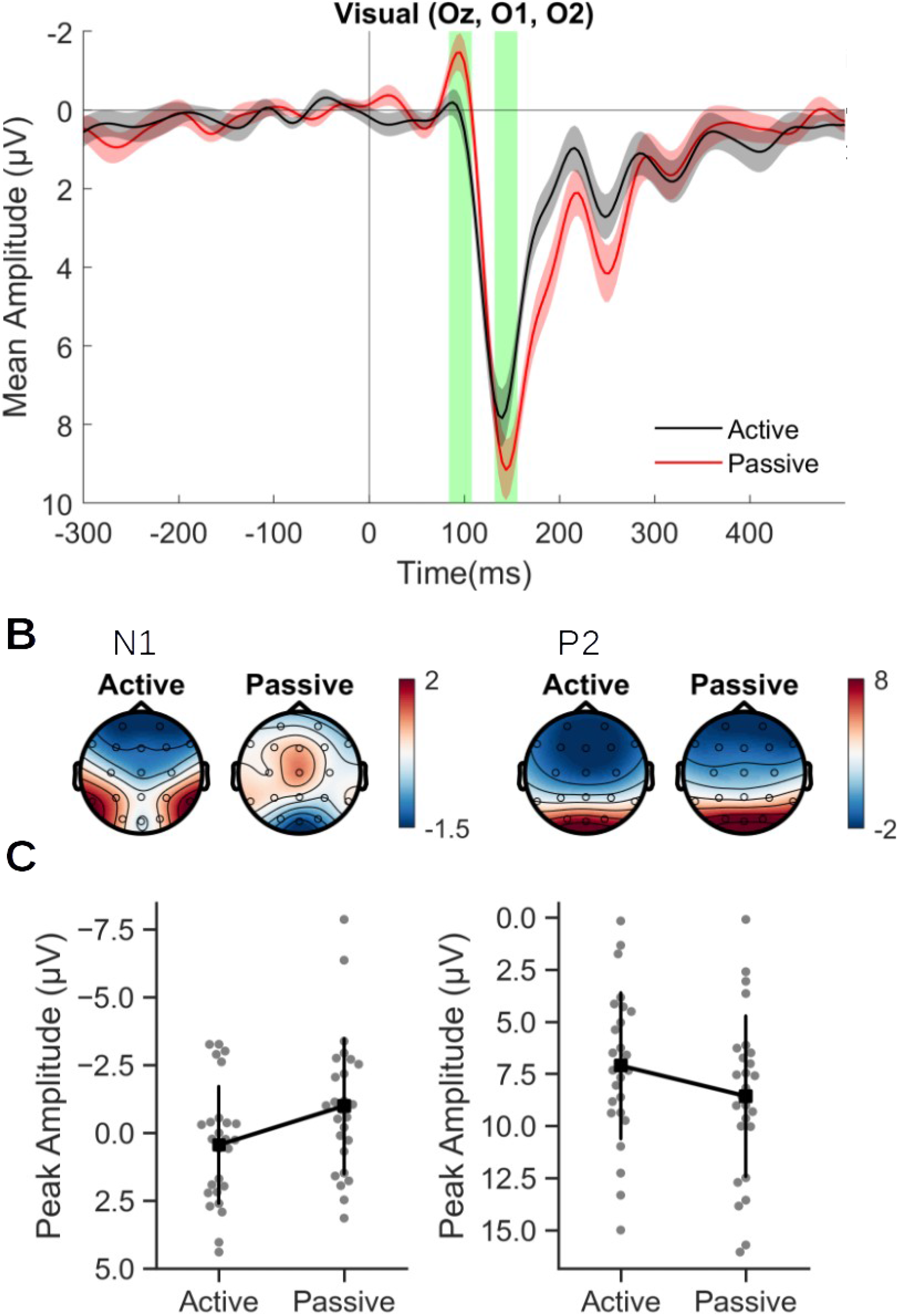
Visual results. (A) Visual ERPs averaged across electrodes Oz, O1 and O2 for active (black) and passive (red) conditions, time locked to the stimulus of interest (0). Red and black shaded areas represent standard deviation. Green shaded areas represent analysis time windows for N1 and P2 peaks. Baseline period −100 to 0ms. (B) Scalp topography maps averaged over the selected time windows for N1 and P2. (C) Mean N1 and P2 amplitudes. Error bars represent standard deviation.

For the auditory condition (Figure 3), the effect action type for N1 was not significant (*t_25_* = −1.87, *p* = .074, *d* = −.37, CI = [−1.195, 0.04]). The effect of action type for P2 was significant (*t_25_* = −2.17, *p* = .04, *d* = −0.43, CI = [−1.648, −0.044]), with more positive values for passive (mean = 2.04) than active movements (mean = 1.9), indicating P2 suppression. However, this effect did not remain significant after removal of the participant whose behavioural data were missing (see Appendix).

**Figure 3.**
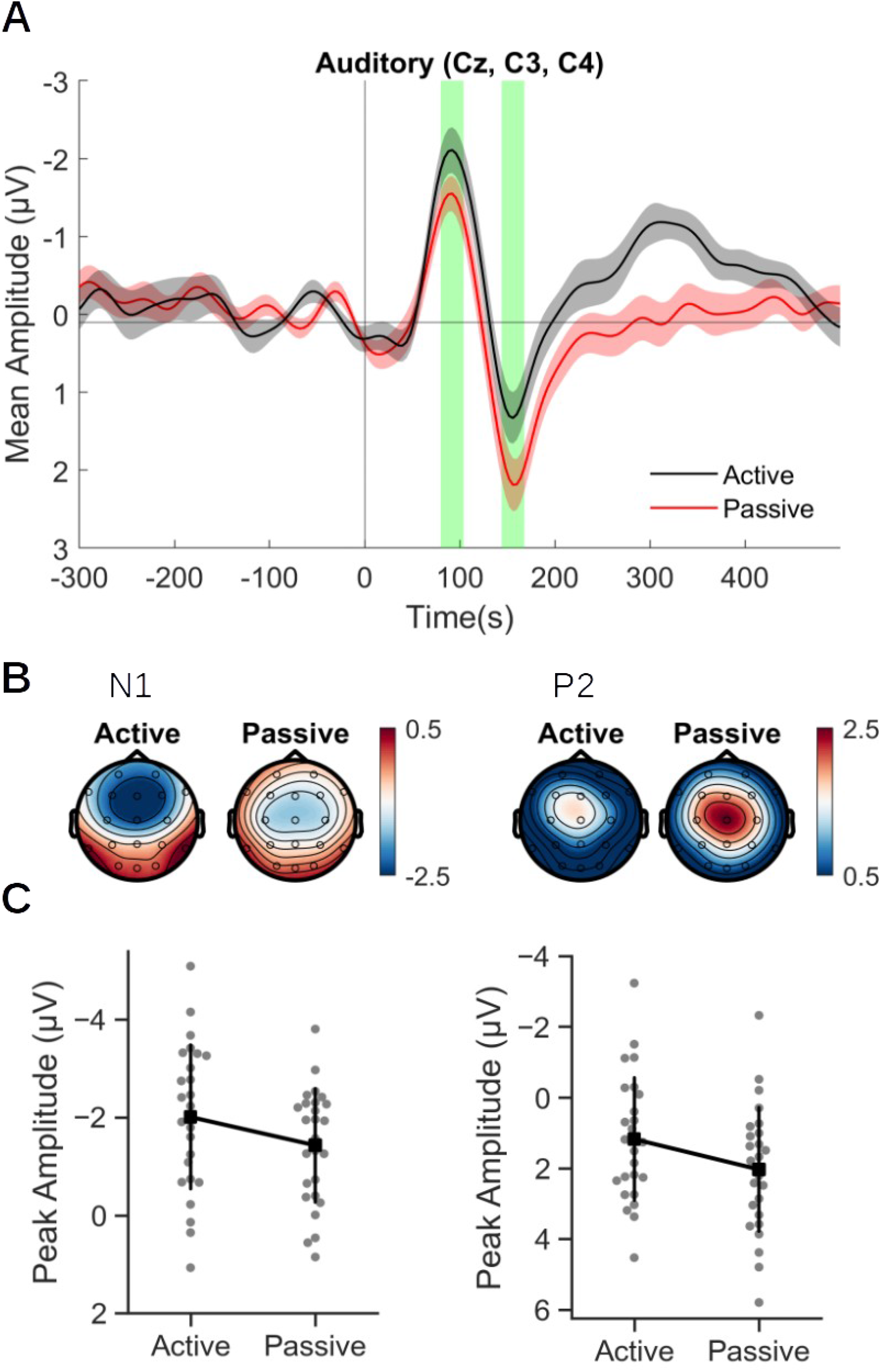
Auditory results. (A) Auditory ERPs averaged across electrodes Cz, C3 and C4 for active (black) and passive (red) conditions, time locked to the stimulus of interest (0). Red and black shaded areas represent standard deviation. Green shaded areas represent analysis time windows for N1 and P2 peaks. Baseline −100 to 0ms. (B) Scalp topography maps averaged over the selected time windows for N1 and P2. (C) Mean N1 and P2 amplitudes. Error bars represent standard deviation.

**Figure 4.**
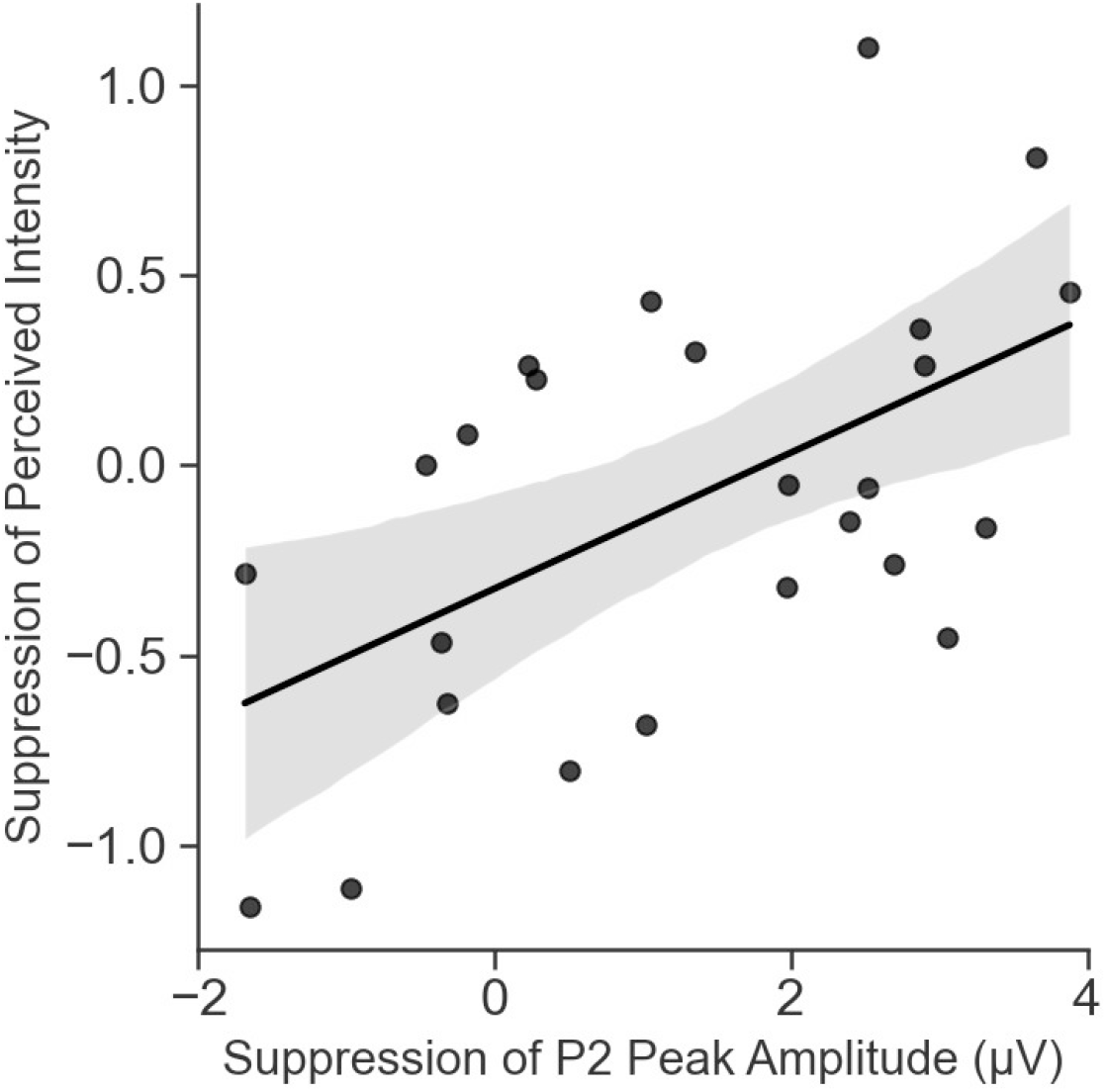
Relationship between suppression of P2 peak amplitude, derived by subtracting active peak values from passive peak values, and suppression of perceived intensity, derived by subtracting active thresholds from passive thresholds.

### 3.3 Correlation

In the visual domain, there was a significant correlation (*r_23_*= .55, *p* = .005) between *P2* suppression (passive – active peak value) and behavioural suppression (active – passive threshold) at the Bonferroni-corrected alpha level of .025. Thus, subjects who judged the intensity of self-generated visual stimuli as darker (lower in intensity) are more likely to also have a reduced P2 amplitude.

The correlation between *N1* suppression (active-passive peak value) and behavioural suppression in the *visual* domain was not significant (*r_23_* = −0.018, *p* = .93). The correlation between *P2* suppression and behavioural suppression in the *auditory* domain was also not significant (*r_23_* = 0.24, *p* = .24).

## 4. Discussion

In this experiment, we investigated neural and behavioural responses to self- and externally-initiated visual and auditory action consequences. We used a PMD to produce involuntary button presses. Using this method, we were able to reduce differences in temporal prediction and tactile sensation between the active and passive conditions. We also included an intensity judgement task to investigate whether neural responses are related to intensity perception.

Our results showed robustly reduced amplitude of N1 and P2 in the primary visual response for self-generated (active) visual stimuli. We observed no significant differences between points of subjective equality between the active and passive conditions in the intensity judgement task in the visual or auditory domains. However, suppression of perceived intensity in the visual intensity judgement task was significantly correlated with suppression of the visual P2 peak. In other words, participants who showed greater P2 suppression in the behavioural task also tended to show greater suppression in the elicited neural activity.

In the visual domain, self-initiated stimuli led to suppressed N1 and P2 components in the primary visual response over the occipital electrodes. This result extends a large body of literature showing sensory suppression of self-produced tones. However, results of previous studies investigating sensory suppression of visual action consequences (Csifcsák et al., 2019; Gentsch & Schütz-Bosbach, 2011; Hughes & Waszak, 2011; Mifsud et al., 2016; Schafer & Marcus, 1973) have provided mixed evidence for sensory suppression in the visual sensory modality. These studies have differed greatly in design, choosing stimuli ranging from low-level flashes (Schafer and Marcus, 1973; Mifsud et al. 2018), arrow shapes (Gentsch & Schütz-Bosbach, 2011) and patterns (Csifcsák et al., 2019; Mifsud, et al., 2016; Hughes & Waszak, 2011) to realistic images such as faces, houses (Hughes and Waszak, 2014), and hands (Csifcsák et al., 2019). Nevertheless, the present results support the notion that efference copy-based (forward model) predictions lead to reduced processing of visual outcomes of actions, reflected in N1 and P2 components.

The majority of previous studies have used some variation of the contingent paradigm. It has been argued that this design could overestimate the effects of sensory suppression or be open to other explanations of the results besides an efference copy-based forward model mechanism (Horváth, 2015). In the present study, we employed an innovative design using a PMD which allowed us to produce involuntary finger movements. Notably, this device allowed for improved control over two factors which are thought to affect sensory attenuation. Firstly, it meant that we were able to make stimuli temporally predictable in both conditions by presenting them at a fixed interval after the button press. This meant that even though the button press itself was not predictable in the passive condition, the resulting stimulus nevertheless was. Secondly, in the classic contingent paradigm, it has been suggested that attention could be drawn to the button press and away from the stimuli in the active condition, leading to the apparent sensory suppression effect. In our design, this is unlikely as the tactile sensation of the finger making contact with the button was present in both conditions. Thus, remaining differences in demand would only reflect processes which are self-initiation (efference copy) related. While controlling these factors, we nevertheless observed robust N1 and P2 suppression in the visual domain and our results therefore corroborate previous sensory suppression studies.

The consequence of voluntary and involuntary movements on initiated stimulation has not been studied extensively in either the auditory or visual domains. Two previous EEG studies have compared electrophysiological responses to tones following voluntary and involuntary movements. Timm et al. (2014) compared responses to sounds triggered by a voluntary button press to ones triggered by similar (but involuntary) movements elicited by TMS over the motor cortex. They observed attenuation of the N1-P2 complex. Similarly, Jack et al. (2021) employed involuntary movements by having the experimenter move the participant’s finger or stimulate the median nerve, producing an involuntary finger movement. They also included semivoluntary conditions in which the participant used a finger from the left hand to push down a finger on the right hand, moving the button and triggering a stimulus, or used the left hand to activate the nerve stimulator placed on their right arm. N1 responses to tones produced by semivoluntary and involuntary movements did not significantly differ from tones which were passively listened to. These results are consistent with forward-model theories of action prediction. They suggest that movement planning, which produces an efference copy of the predicted sensory action consequences is necessary to produce neural sensory suppression in the auditory sensory modality. Our results complement these findings by suggesting that movement planning is also necessary for the neural suppression of visual action consequences.

As previously discussed, suppression of auditory N1 and P2 to self-initiated tones has been shown in a substantial number of previous studies (Bäß et al., 2008; Martikainen et al., 2005; Mifsud et al. 2016; SanMiguel et al., 2013). Therefore it is unusual that we did not observe this in our data. Instead, in the auditory condition, we observed no significant differences in the N1 time window. Furthermore, while the P2 did show the expected effect direction, the result did not remain significant after the removal of the one participant whose behavioural data were not available (see Appendix). This suggests that the results for the auditory condition were not robust. The results are also inconsistent with the visual condition, where we observed a large sensory suppression effect. We suspect that this unexpected result may have been caused by unavoidable noise produced by the button mechanism in the passive condition. While we did attempt to control this factor using pink noise throughout the experiment, it is possible that the noise from the button caused a sensory gating effect in the passive condition, causing the stimulus-locked ERP to be smaller than expected. Further evidence for this hypothesis is provided in the appendix.

The main results of the intensity judgement task did not show significant differences in judgements of intensity when comparing stimuli produced by active and passive button presses. This result stands in contrast to Lubinus et al. (2021) who found that self-initiated visual stimuli were perceived as significantly darker than externally-generated stimuli in a similar intensity judgement task. However, they presented their stimuli for 1 second, whereas we presented ours for only 50 milliseconds. We chose this presentation length to be consistent with previous EEG studies (e.g. Bäß et al., 2008, Timm et al., 2014; Timm et al., 2016). However, it is possible that this short duration affected the judgement of the stimuli. As previously discussed, EEG studies of visual sensory suppression have employed a wide range of different stimuli. Likewise, behavioural measures were different between studies, with previous tasks including intensity judgement (Lubinus et al., 2021; Yon & Press, 2017), speed judgement (Dewey and Carr, 2013), contrast discrimination (Roussel et al., 2013, Vasser et al., 2019), stimulus detection (Cardoso-Leite et al., 2010; Schwarz et al., 2018), delay detection (Arikan et al., 2019; Kavroulakis et al., 2021; Pazen et al., 2020; Schmitter et al., 2021; Straube et al., 2017; Uhlmann et al., 2020; Uhlmann et al., 2021; van Kemenade et al., 2016; van Kemenade et al., 2019) and similtanaiety judgement (Arikan et al., 2017). Furthermore, Mifsud et al. (2018) have shown that the causal likelihood of a visual stimulus occurring with a specific motor action can affect sensory attenuation. They demonstrated that saccade-initiated flashes elicited greater sensory attenuation than button press-initiated flashes. Therefore the precise conditions necessary to produce robust sensory suppression in the visual domain remain unclear. Nevertheless, our task captured sensory suppression at least to some degree, but results seem to be covered by large individual differences as indicated by the correlation analyses (see next paragraph). Our study demonstrates the relevance of the inclusion of behavioural measures as neural responses might not directly reflect the perceptual level, leading to false conclusions about perceptual suppression based on neural responses.

As one of our key findings, the correlation between suppression in the visual intensity judgement task and suppression of P2 peak amplitudes is the first evidence to show a link between electrophysiological suppression and perception of stimulus intensity. Participants that showed greater levels of suppression in the intensity judgement task, also showed greater suppression of the P2 component of the visual event-related potential. Individual differences in neural suppression related to behaviour and been previously observed. For example, in an fMRI study (Schmitter et al., 2021) investigating the neural processing of continuous and discrete action outcomes, suppression on the behavioural level in a delay detection task was negatively correlated with BOLD suppression (passive - active) in left middle occipital gyrus specifically for discrete action outcomes. Despite the observed correlation and previous reports, it is unclear which factors could be responsible for these individual differences. Subclinical symptoms of psychosis could represent one possibility. The ‘continuum of psychosis’ is a theory which suggests that psychotic-like experiences may be distributed along a continuum ranging from low severity in the healthy population to high severity in individuals with psychotic disorders, such as Schizophrenia (Verdoux & van Os, 2002). It has been suggested that positive symptoms of Schizophrenia could be explained by an underlying deficit in efference copy-based forward model mechanisms (Pynn & DeSouza, 2013). Evidence for this is shown in previous studies which find reduced sensory suppression of self-generated sensory action consequences in patients with Schizophrenia (Ford, Gray, et al., 2007; Ford, Roach, et al., 2007 Ford et al., 2014). Furthermore, individuals scoring higher on Schizotypy, a personality trait which describes psychotic-like experiences in non-clinical individuals, show lower levels of sensory suppression of self-generated speech (Oestreich et al., 2015). Therefore, the individual differences here could also be related to variation in schizotypal personality. In any case, the visual P2 may be behaviourally relevant. On the other hand, our results challenge previous studies that interpreted their findings of N1 suppression as an indicator of suppression on a perceptual level. Future studies should aim to establish these relationships further by investigating whether individual personality differences are related to sensory suppression in the visual domain. Further work also needs to be done to establish the electrophysiological correlates of sensory suppression in other behavioural tasks where action prediction effects are found.

## 5. Conclusions

The aim of this study was to investigate action-related sensory suppression during self- and externally-generated visual and auditory feedback. Using a novel version of the typical paradigm, we included involuntary passive movements which allowed us to control tactile sensory feedback and temporal prediction of stimuli. We showed that self-generated compared to externally-generated visual action consequences elicited reduced activity in the primary visual response over the occipital electrodes. Furthermore, we demonstrated for the first time that participants who showed greater suppression in the perception of visual stimulus intensity also showed greater suppression in the elicited neural activity. Taken together, these results support the idea of an efference copy-based forward model mechanism in the visual sensory modality. They also speak for the role of movement planning in visual sensory suppression. Finally, they suggest that early components of the primary visual response are behaviourally relevant in perceiving the intensity of self-generated visual action consequences.

## Author Note

This work was funded by the German Research Foundation, Germany, through the IRTG 1901 “The Brain in Action” (DFG GRK 1901/2), the SFB/TRR 135 (project no. 222641018, project A3) and is supported by “The Adaptive Mind”, funded by the Excellence Program of the Hessian Ministry for Science and the Arts (to BS, TK). BS is funded by the DFG (STR 1146/15-1 Grant Number 429442932, STR 1146/9-1/2, Grant Number 286893149). YH is funded by the DFG (HE8029/2-1). The funders had no role in the study design, collection or analysis of data, writing or the decision to publish the article. We thank Katharina Schuster and Mona Rauschkolb for assistance with data collection and translation and Ruslan Spartakov and Julia Oppermann for assistance with data collection.

## Author Credit Statement

Conceptualization: EO, BS, YH, TK

Methodology: EO, YH, BS

Software: EO

Validation: EO

Formal analysis: EO, YH

Investigation: EO

Resources: BS, TK

Data curation: EO

Writing – Original Draft: EO

Writing – Review & Editing: BS, YH, TK

Visualisation: EO

Supervision: BS, YH, TK

Project administration: BS, TK

Funding acquisition: BS, TK

## Data and Code Availability Statement

Group-level data and accompanying scripts supporting the conclusion presented in this study are available from OSF: DOI 10.17605/OSF.IO/XGQP7.

## Appendix. Supplementary Materials

### Auditory Condition Confound

There was a discrepancy between the visual and auditory results. As hypothesised, the visual results showed a reduction in amplitude in the ERP elicited by self-generated visual stimuli. However, no significant differences between active and passive were observed for N1 in the auditory condition. Furthermore, the P2 effect was not robust, no longer showing a significant effect after the removal of one participant whose behavioural data were missing. We suspected this being due to unavoidable noise caused by the electromagnet mechanism in the PMD, despite our playing pink noise over earphones. To verify this hypothesis, we examined event-related potentials time-locked to the button press. Data were preprocessed as described in the methods. Data were then segmented from 400ms before to 400ms after the button press and from 400ms before to 400ms after the trial onset. The button press segments were baseline corrected to the 200ms preceding the trial onset. ERPs were derived by averaging across participants and electrodes Cz, C3 and C4.

Figure S1 shows ERPs for the auditory, visual and control conditions, time-locked to the button press. Passive conditions show a negative peak, on average 74.7 ms after the button press, in frontocentral electrodes. Given that the location and latency of this peak are typical of an auditory N1 and the fact that it is only present in the passive condition, it is likely that the peak is related to the button noise. In sensory gating paradigms, in which a pair of tones or clicks is delivered with a short interval in between, auditory N1 components are reduced after the second sound (e.g. Boutros et al., 2004; Brockhaus-Dumke et al., 2008). In our paradigm, in the passive condition, uncontrollable noise from the button was followed shortly by the intended tone stimulus. This could have resulted in a sensory gating effect, causing a lower amplitude in the passive stimulus-locked ERP, which could explain the unexpected effect direction seen in this condition.

**Figure S1.**
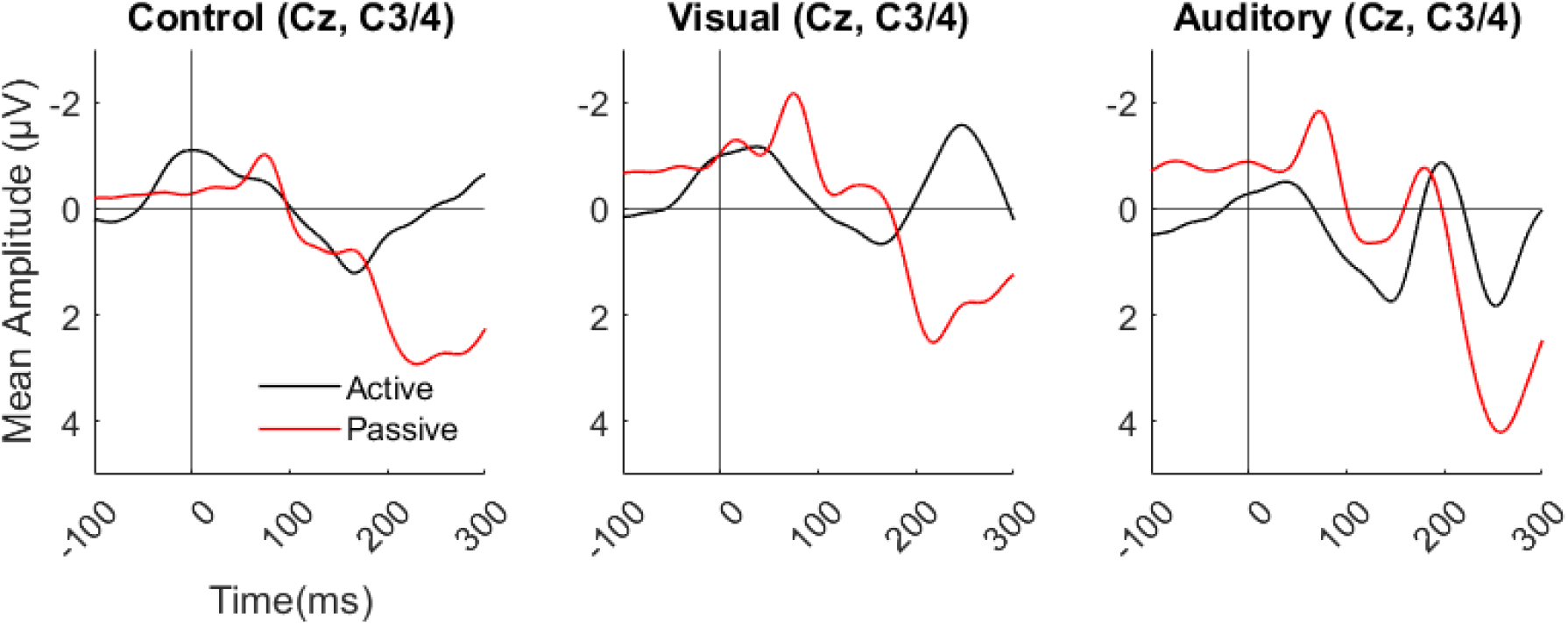
Average ERPs at C3/4 and Cz, time locked to button press (0) for control, visual and auditory conditions.

### Additional Analysis

The behavioural data belonging to one participant were missing due to a technical error. To verify that the inclusion of this participant’s EEG data did not affect the results, we repeated all ERP analyses without this participant’s data included.

For visual conditions, the effect of action type for N1 (*t_24_*= 4.34, *p* < .001, *d* = 0.87, CI = [0.781, 2.197]) and P2 (*t_24_*= −3.87, *p* < .001, *d* = −0.77, CI = [−1.996, - 0.608]) both remained significant at the *p* < .05 level.

For auditory conditions, the effect of action type for N1 remained non-significant at the *p* < .05 level (*t_24_* = −1.99, *p* = .058, *d* = −0.4, CI = [−1.265, 0.003]). However, the effect of action type for P2 was no longer significant (*t_24_* = −1.86, *p* = .075, *d* = −.37,CI = [−1.47, 0.076]).

